# The seed mitochondrial proteome of *Lupinus albus* provides insight into energy metabolism during germination

**DOI:** 10.1101/2025.09.19.677279

**Authors:** Cecile Angermann, Hans-Peter Braun, Tatjana M. Hildebrandt

## Abstract

Despite their essential role in fueling the onset of metabolism, the composition and functional state of seed mitochondria are still a matter of debate. Using white lupin (*Lupinus albus*) as a model, we provide a comprehensive proteomic characterization of seed mitochondria in a legume with protein-rich reserves. Highly enriched mitochondrial fractions isolated from quiescent seeds revealed fully assembled oxidative phosphorylation (OXPHOS) complexes and supercomplexes, demonstrating that mitochondria are preconfigured for immediate energy production upon imbibition. Quantitative proteomics identified 1,162 mitochondrial proteins and highlighted marked tissue-specific differences compared to leaves, including distinct transporter profiles and a dehydrogenase complement supporting amino acid and fatty acid catabolism. Early germination was accompanied by remodeling of coenzyme metabolism and transport capacity, while core respiratory complexes remained stable. Notably, ∼12% of the proteome consisted of uncharacterized proteins, many of which displayed dynamic changes during early germination, suggesting yet undiscovered, potentially legume-specific mitochondrial functions.

## Introduction

Mitochondria are central integrators of energy metabolism in plants. They generate ATP through the combined action of the tricarboxylic acid (TCA) cycle, other catabolic pathways and the respiratory chain, providing an essential complement to photosynthesis, especially in darkness or in non-green tissues. Beyond energy supply, mitochondria are also key players in redox balance, signal transduction, programmed cell death, and the coordination of metabolic networks (Møller et al., 2021). The complexity of mitochondrial function is reflected in their molecular and structural organization. Plant mitochondria are small (1–3 µm) double-membrane organelles with cristae and a protein-rich matrix (Logan, 2006; Taiz et al., 2023). They are highly dynamic, moving along the cytoskeleton and undergoing fusion and fission (Logan, 2006; 2010). Their endosymbiotic origin is evident from the presence of their own genome (Brennicke and Leaver, 2007). Compared with animals, plant mitochondrial genomes are strikingly larger (10–40-fold), but gene-sparse and dominated by repeats (Gualberto and Newton, 2017; Alverson et al., 2011). Only 1–2% of the proteome is mitochondrially encoded, while the vast majority of the ∼2,000–3,000 proteins are nuclear-encoded and imported via N-terminal targeting sequences (White and Scandalios, 1989; Rao et al., 2017). The dense protein environment of the matrix promotes the formation of metabolons, which are functional enzyme complexes that enable the direct channelling of metabolites (Sweetlove and Fernie, 2013). This increases the efficiency of metabolic processes. They also contribute to a high buffer capacity, stabilising the mitochondrial redox balance and preventing toxic increases in coenzyme concentrations (Kasimova et al., 2006).

Technological advances in isolation and analysis methods such as tag-based pull-down techniques now allow mitochondria to be isolated rapidly and specifically, even from individual tissues and developmental stages (Boussardon et al., 2020; Kuhnert et al., 2020; Niehaus et al., 2020). However, mitochondrial preparations from low abundant material remains a challenge (Ditz et al., 2025; Boussardon et al., 2025). Current studies suggest that mitochondria can be considered plastic in terms of their number, shape and function. These properties vary depending not only on the cell type (Logan and Leaver, 2000), but also on the developmental stage of the plant (Paszkiewicz et al., 2017; Rodríguez et al., 2015).

Seeds provide a particularly striking example of a tissue requiring high metabolic plasticity. They tolerate extreme desiccation, remaining viable at water contents as low as 5-15% for years or even decades and thus enable long-term survival and dispersal of plants, even under harsh or fluctuating environments (Bewley et al., 2013 Linkies et al., 2010). This remarkable resilience depends on a suite of protective mechanisms, including accumulation of osmolytes, non-reducing sugars, antioxidant systems, and specialized proteins such as LEA (Late Embryogenesis Abundant) proteins and heat shock proteins that stabilize cellular structures during drying (Sano et al., 2016). Upon rehydration under favourable environmental conditions, metabolic activity resumes within minutes (Paszkiewicz et al., 2017; Nietzel et al., 2020). Stored carbon and nitrogen reserves are remobilized to provide precursors and energy for repair of storage-induced cellular damage, the resumption of transcription and translation, and the synthesis of metabolites and cellular structures to support cell expansion and growth (Weitbrecht et al., 2011; Paszkiewicz et al., 2017; Law et al., 2012). Mitochondria provide ATP efficiently through respiration and are thus particularly important for seed vigor with impaired mitochondrial function frequently leading to germination defects (Ferguson et al., 1990; Kühn et al., 2015; Racca et al., 2022).

Despite their central role, the nature of mitochondria in dry seeds remains debated. Early electron microscopy studies described poorly differentiated “promitochondria” lacking cristae and respiratory chain complexes in different plant species (Attucci et al., 1991; Logan et al., 2001; Howell et al., 2006). In this model, mitochondria would only achieve full functionality after imbibition through rapid synthesis and assembly of oxidative phosphorylation (OXPHOS) complexes (Law et al., 2012; Paszkiewicz et al., 2017). More recent proteomic and functional studies in *Arabidopsis thaliana*, however, argue that seed mitochondria are already competent, containing pre-assembled respiratory complexes and unique protein adaptations for desiccation tolerance (Nietzel et al., 2020; Ditz et al., 2025). This unresolved discrepancy highlights the need for in-depth characterization of mitochondrial proteomes across diverse seed types.

In this study, we characterize the mitochondrial proteome of white lupin (*Lupinus albus*), a legume of agronomic and ecological importance. Lupin seeds are notable for storing a large fraction of their reserves as protein (∼40%), making them an ideal system to study seed-specific adaptations in amino acid and energy metabolism (Angermann et al., 2024). Their large size facilitates experimental handling and yields sufficient material for high-purity organelle preparations. Leveraging our curated *L. albus* proteome annotation database, we present a comprehensive analysis of seed mitochondria, revealing distinct compositional features that reflect their specialized role in seed physiology.

## Results

### Highly enriched mitochondrial preparations from stratified white lupin seeds provide insight into the seed mitochondrial proteome composition

Upon imbibition under favorable conditions, white lupin seeds rapidly initiate respiration and complete germination with radicle emergence after approximately 30 h (Fig. 1, Angermann et al., 2024). This instant metabolic activation suggests that mitochondria are preconfigured to support the transition from quiescence to active growth. Yet, the mechanisms by which mitochondria are primed for this role, and the molecular features that enable them to drive seed metabolism in the context of protein-rich reserves, remain poorly understood. To address these questions, we aimed to isolate highly enriched mitochondrial fractions from seeds in a quiescent state prior to germination. Direct isolation from dry seeds proved technically challenging, prompting us to apply a cold stratification step. Lupin seeds soaked in demineralized water for 24h at 4°C underwent water uptake and swelling while maintaining minimal metabolic activity (Fig. 1). Following transfer to room temperature, they remained viable and proceeded to germination, confirming that the treatment preserved seed physiology.

**Fig. 1:**
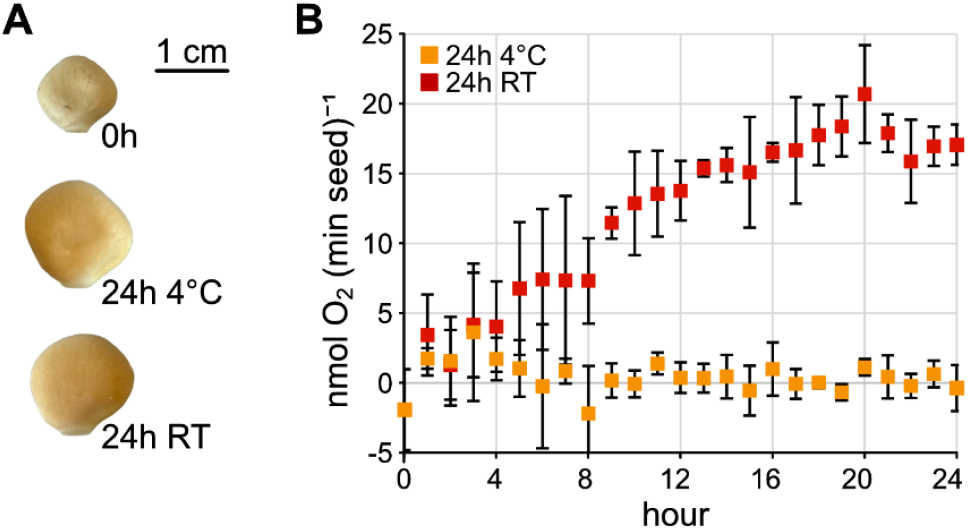
Imbibed *Lupinus albus* seeds rapidly start respiring at room temperature but not in the cold. (**A**) Pictures of representative *Lupinus albus* seeds in the dry state (0h) as well as after 24h imbibition at 4°C (24h 4°C) or room temperature (24h RT) (**B**) Respiration rates of *Lupinus albus* seeds during imbibition at 4°C and at room temperature (RT). Data presented are means ± SD (n = 3).

Proteome analysis by two-dimensional (2D) blue-native/SDS polyacrylamide gel electrophoresis (BN/SDS-PAGE) revealed that mitochondria from lupin seeds contain fully assembled OXPHOS complexes and supercomplexes (Fig. 2A). Although the amount of seed tissue was equal for all mitochondrial isolations, the preparations from dry seeds showed only faint bands and spots, most likely reflecting inefficient recovery of the organelles from this starting material. In contrast, 2D gels from imbibed seeds clearly showed the characteristic spot pattern of the respiratory chain complexes I-IV, the ATP synthase (complex V), the supercomplex composed of complex I and dimeric complex III and further complexes like the HSP60 complex, all well known from work with *Arabidopsis thaliana* (Klodmann et al., 2011).

**Fig. 2:**
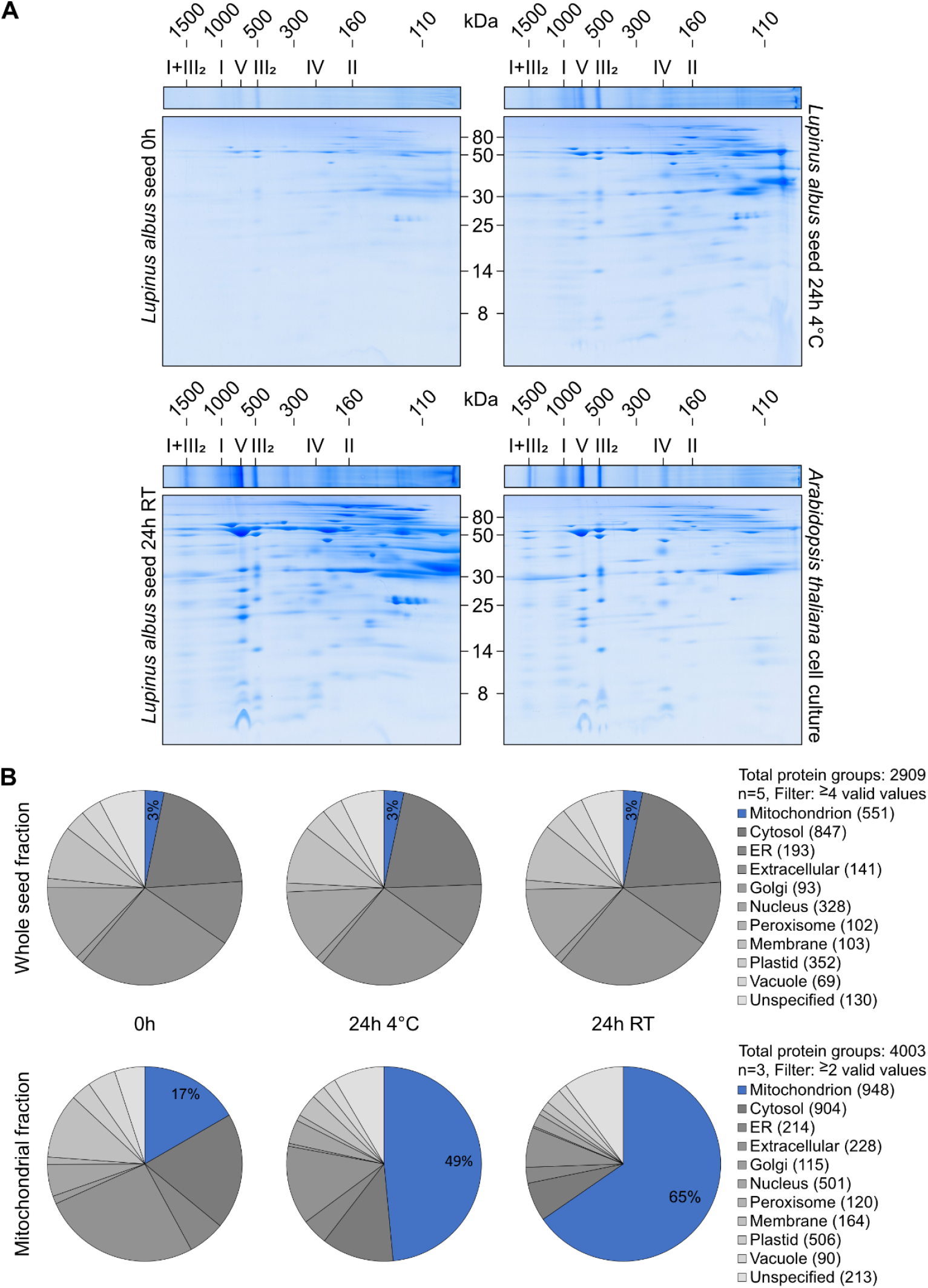
*Lupinus albus* seed mitochondrial proteomics. (**A**) Analyses of mitochondrial protein complexes from *Lupinus albus* (0h: top left; 24h 4°C: top right; 24h RT: bottom left) and *Arabidopsis thaliana* (cell culture: bottom right) by 2D BN/SDS-PAGE. Lanes of 1D BN gels were transferred horizontally onto SDS gels for electrophoresis in orthogonal direction (see “Materials and methods” section for details). The identities of the protein complexes as well as molecular masses [kDa] are indicated above the gels (identifications based on *A. thaliana* reference gels, see Klodmann et al., 2011). Molecular masses of monomeric standard proteins [kDa] are indicated between the gels. (**B**) Quantitative proteome composition of total seed extracts (top) and isolated mitochondrial fractions (bottom). Proteins were assigned to subcellular compartments based on SUBA5 (Hooper et al., 2014) and mass fractions [%] were calculated from cumulated iBAQ values for all compartments. Numbers in parentheses indicate the number of different distinct proteins detected in the respective category. Data presented are means of five replicates for whole seed samples and means of three replicates for mitochondrial fractions. The complete dataset is provided as Supplementary Dataset S1.

We analyzed the proteome composition of the enriched mitochondrial fractions and the original protein extracts from seeds by LC-MS/MS (Supplementary Dataset S1). For protein annotation we used our manually curated extended *Lupinus albus* annotation database (Angermann et al., 2024), which based on sequence alignments also provides a link to the extensive resources available for *Arabidopsis thaliana* such as functional annotation of pathways according to the MapMan system (https://mapman.gabipd.org; Thimm et al., 2004) and assignments to subcellular compartments using the SUBA5 database (Hooper et al., 2014; Hooper et al., 2017; Hooper et al., 2022). Based on SUBA5, 551 distinct proteins were assigned to mitochondria in total seed extracts and 948 distinct proteins in the mitochondrial fractions (Supplementary Dataset S1). We calculated the quantitative composition of the individual proteome samples based on normalized iBAQ values to estimate the purity of the mitochondrial preparations (Fig. 2B, Supplementary Dataset S1). Mitochondrial proteins accounted for 3% of the total seed protein content. Their proportion increased to 17% in the mitochondrial fractions prepared from dry seeds (6-fold enrichment), and to 49% and 65% in mitochondrial fractions obtained from seeds imbibed at 4°C and room temperature (16- and 20-fold enrichment, Fig. 2B). Thus, cold stratification proved effective in facilitating mitochondrial isolation from metabolically inactive seeds, enabling the preparation of enriched mitochondrial fractions suitable for proteomic analysis prior to the onset of germination.

### Identification of new mitochondrial proteins in white lupin seeds

Proteins with a mitochondrial annotation according to SUBA5 were highly enriched in the mitochondrial fractions prepared from *L. albus* seeds imbibed for 24h at 4°C or RT compared to the total seed extract reflected by their strongly and significantly increased relative abundance in the mitochondrial preparations (Fig. 3A, dark blue dots, Supplementary Dataset S2). To identify previously unknown *bona fide* mitochondrial proteins we defined thresholds for log2-fold change (mitochondrial fraction/total protein fraction) and −log10 p-value above which 70% of all proteins were annotated as “mitochondrial” (Fig. 3A, dotted lines, Supplementary Dataset S2, Supplementary Fig. S1). From the 175 proteins above this threshold without a mitochondrial SUBA annotation, 29 could be assigned to mitochondria based on complementary evidence (light blue), 84 proteins could not be confidently linked to an Arabidopsis homolog and were thus lacking a localization annotation (orange), while 62 proteins remained as strong candidates for novel mitochondrial components (red).

**Fig. 3:**
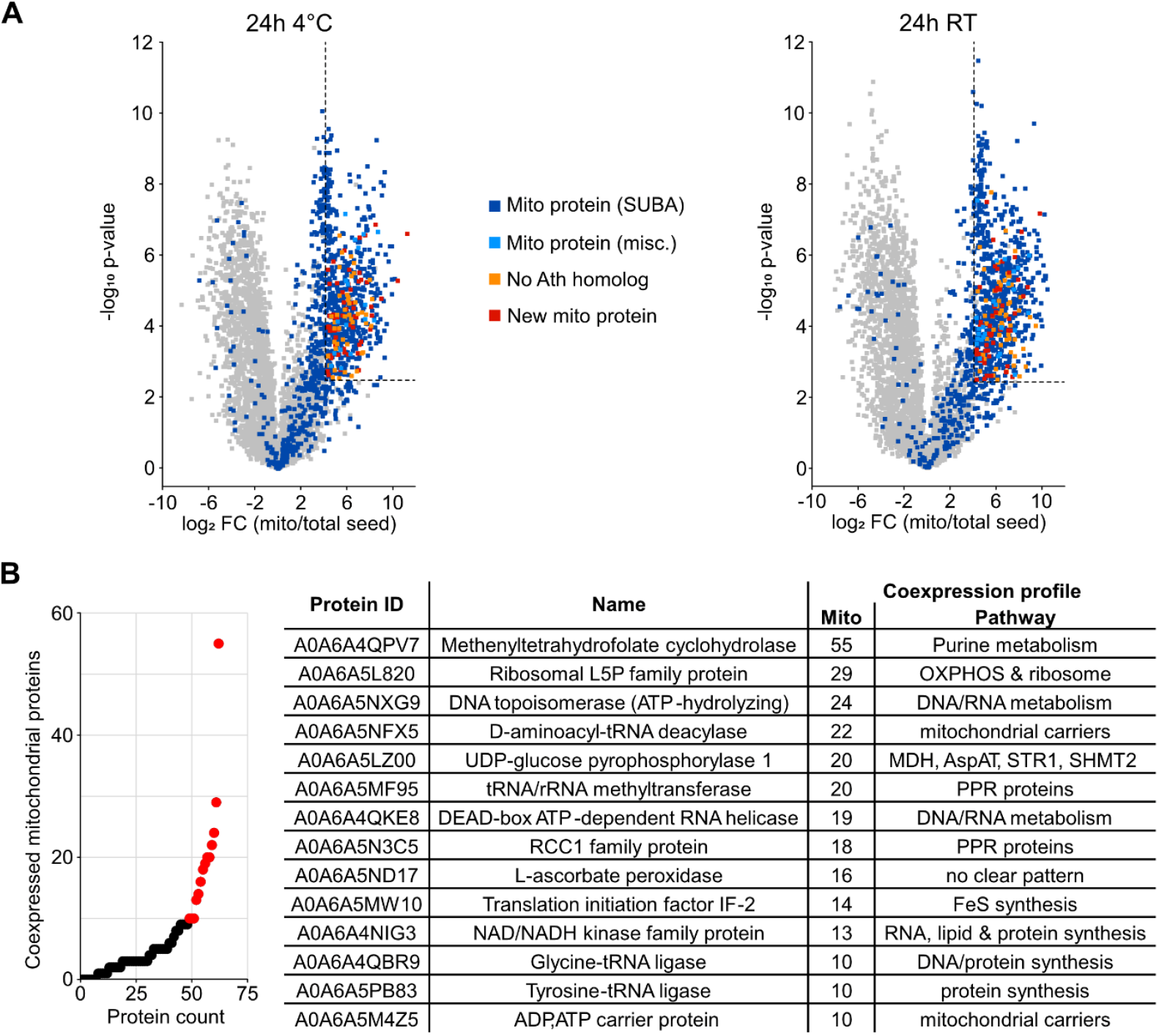
Identification of new seed mitochondrial proteins. **(A)** Volcano plots illustrating differences in relative protein abundances in mitochondrial fractions compared to total seed extracts. Dark blue: proteins annotated as mitochondrial in SUBA5, light blue: proteins annotated as mitochondrial based on other sources (MULocDeep, Jiang et al., 2021, Jiang et al., 2023), orange: no prediction possible due to missing Arabidopsis homolog, red: new mitochondrial protein candidate predicted as non-mitochondrially localized in *Arabidopsis thaliana*, but according to our enrichment analysis located in mitochondria in *Lupinus albus*. The dotted line indicates the threshold for new mitochondrial protein candidates. (**B**) Coexpression analysis for new mitochondrial protein candidates: The graph illustrates the fraction of genes encoding mitochondrially localized proteins in the 100 most strongly coexpressed genes for each candidate (ATTED-II; Obayashi et al., 2018). Additional information is given in the table for the candidates with the highest mitochondrial association (red dots). The complete dataset is provided as Supplementary Datasets S2 and S3.

To gain further insight into the functional context of these candidates, we performed coexpression analyses of their A. thaliana homologs using ATTED-II (Obayashi et al., 2018, Supplementary Dataset S3). Fourteen candidates displayed coexpression patterns in which at least 10% of the top 100 co-expressed genes were mitochondrially localized (Fig. 3B). These coexpression profiles suggest links to processes such as mitochondrial translation, RNA editing, and transcription, as well as metabolite transport, energy metabolism, and amino acid metabolism. Such associations highlight these proteins as promising targets for further investigation into mitochondrial function in seeds.

### The quantitative proteome composition of white lupin seed mitochondria

Including the newly identified candidates our *L. albus* seed mitochondrial proteome dataset comprises 1,162 distinct proteins (Supplementary Dataset S4). For functional annotation, we used a modified and manually curated version of the MapMan system (Thimm et al., 2004), which divides the *L. albus* proteome into 20 major categories with two levels of sub-categories (Angermann et al., 2024). The quantitative distribution of protein abundances across functional categories is illustrated by Proteomaps (Liebmeister et al., 2014, Fig. 4A, Supplementary Dataset S4). As expected, oxidative phosphorylation (OXPHOS) dominated the seed mitochondrial proteome, with respiratory chain complexes representing ∼40% of the total protein mass. ATP synthase subunits α and β were the most abundant individual proteins (2.5% each), and complex V as a whole accounted for 17% of mitochondrial protein mass. Voltage-dependent anion channel (VDAC) isoforms were also highly abundant, collectively comprising 7% of the proteome and dominating the functional category “transport” (11%). The top 10 most abundant seed mitochondrial proteins also include an NAD-dependent formate dehydrogenase, a cystationine beta-synthase family protein, an adenine nucleotide translocator, malate dehydrogenase, and a homolog of *A. thaliana* early nodulin 93 (Supplementary Dataset. S4A, sorted by rank: column BH). A striking feature of the *L. albus* seed mitochondrial proteome is the large fraction of uncharacterized proteins consisting of 238 distinct proteins covering ∼12% of the mitochondrial protein mass (Supplementary Dataset. S4B “Not assigned”). For 102 of these, reliable functional annotation based on *A. thaliana* was not possible due to only weak homology, suggesting they may represent lupin- or legume-specific mitochondrial proteins.

**Fig. 4:**
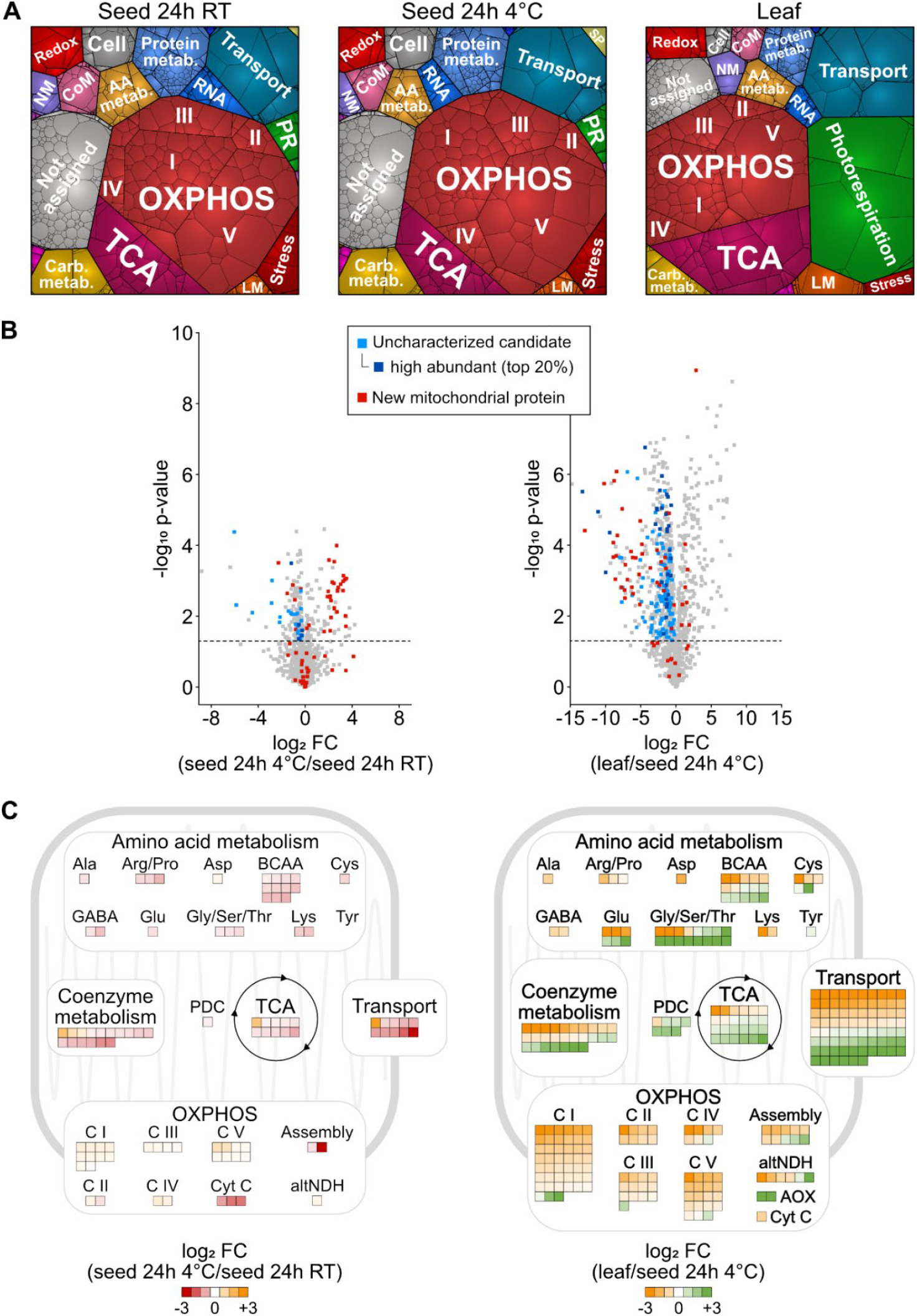
Comparative analysis of mitochondrial proteomes in seeds and leaves. To ensure comparability across datasets with different enrichment levels, all proteomes were filtered for mitochondrial proteins (SUBA5 annotations plus the new seed mitochondrial proteins identified in this study) before normalization and statistical analysis (Supplementary Dataset S4A). **(A**) Proteomaps illustrating the quantitative composition of the mitochondrial proteome of seeds imbibed for 24h at 4°C or at RT and of leaves. Proteins involved in similar cellular functions are arranged in adjacent locations and visualized by colors. Proteins are shown as polygons whose sizes represent their relative abundance (%iBAQ). Proteomaps were produced using the tool provided at https://bionic-vis.biologie.uni-greifswald.de/ (Liebermeister et al., 2014). Data presented are means of three biological replicates. (**B**) Volcano plots illustrating log_2_-fold changes (FC) in relative protein abundances between the mitochondrial proteome of seeds imbibed for 24h at 4°C and seeds imbibed for 24h at RT (left) or leaves and seeds imbibed for 24h at 4°C (right). Dashed lines mark the significance threshold. The new mitochondrial proteins identified in this study are highlighted in red. Uncharacterized proteins significantly induced during germination (left) or in seeds vs. leaves (right) are highlighted in blue with dark blue representing high abundant proteins (top 20%). Data presented are means of three biological replicates. (**C**) Relative abundance of proteins involved in major mitochondrial pathways in the mitochondrial proteome of seeds imbibed for 24h at 4°C vs. seeds imbibed for 24h at RT (left) or leaves vs. seeds imbibed for 24h at 4°C (right). The colored squares represent significant log_2_-fold changes in the abundance of individual enzymes and their subunits (n = 3) with orange indicating increased abundance in quiescent seeds (24h at 4°C) in both comparisons. AA metab.: amino acid metabolism; altNDH: alternative NADH dehydrogenase; BCAA: branched chain amino acid; C I: Complex I; C II: complex II; C III: complex III; C IV: complex IV; C V: complex V; CoM: coenzyme metabolism; Cyt C: cytochrome C; LM: lipid metabolism; NM: nucleotide metabolism; PDH: pyruvate dehydrogenase; PR: photorespiration; SP: storage protein.

### Seed mitochondrial proteome remodelling during the transition to metabolic activity

To capture initial changes associated with germination, we compared mitochondrial fractions from seeds imbibed for 24h at 4°C (hydrated but metabolically inactive) with those from seeds imbibed for 24h at room temperature (actively germinating). The proteomaps indicate substantial stability of the core proteome (Fig. 4A), consistent with the presence of fully assembled respiratory complexes already in dry seeds as revealed by 2D BN/SDS-PAGE (Fig. 2A). Statistical analyses identified 158 proteins that increased and 108 that decreased significantly during the first 24h of germination (Fig. 4B, Supplementary Dataset S4A, filter column G). The strongest responses were detected in upstream supply and support pathways of oxidative phosphorylation (Fig. 4C, Supplementary Dataset S4A, filter column BC). Enzymes of the TCA cycle and amino acid catabolism as well as malic enzyme increased in abundance. Several metabolite and protein transporters, including the dicarboxylate carrier and TIM subunits, were strongly induced. Coenzyme metabolism was also upregulated, with elevated levels of enzymes involved in lipoic acid and thiamine biosynthesis and Fe–S cluster assembly (Fig. 4C, Supplementary Dataset S4A, filter column BC).

Notably, several of the seed mitochondrial proteins strongly induced during the transition to metabolic activity remain uncharacterized (Fig. 4B blue dots, Supplementary Dataset S4A, filter column BD). 17 of the 31 uncharacterized germination induced proteins had only weak homology to Arabidopsis genes, suggesting lupin- or legume-specific functions. Coexpression analysis of the remaining candidates indicated potential roles in ribosome biogenesis and cofactor biosynthesis (Supplementary Dataset S5). Among the new mitochondrial proteins identified in this study, six increased during germination whereas 29 decreased, suggesting that they may play specific roles in dry seed mitochondria that are lost as metabolism is reactivated.

### Distinct features of the seed mitochondrial proteome

To assess tissue specificity of the mitochondrial proteome, we compared seed mitochondria with those from *L. albus* leaves. Leaf mitochondria were less challenging to isolate and yielded highly enriched fractions, with 89% of protein mass annotated as mitochondrial (Supplementary Dataset S1). Visualization of the quantitative proteome compositions using proteomaps immediately highlights clear tissue-specific differences, both in highly abundant mitochondrial proteins and in the representation of major pathways (Fig. 4A). Statistical analyses further resolved the pronounced compositional differences reflecting distinct metabolic requirements of the two tissues (Fig. 4B, C, Supplementary Dataset S4A). A total of 566 mitochondrial proteins were significantly more abundant in seeds than in leaves (Supplementary Dataset S4A filter column H). Pathway enrichment analysis revealed a strong overrepresentation of OXPHOS proteins, particularly subunits of complex I, within this group (Supplementary Dataset S4C). In the 227 proteins with a significantly higher abundance in leaves, TCA cycle enzymes were enriched as well as enzymes involved in photorespiration and transport. Transporter profiles differed strikingly between the two tissues. Of the 101 identified transporters and protein translocators, 76 showed significant changes in abundance, indicating contrasting requirements for respiratory substrates, metabolite exchange, and protein import (Fig. 4C, Supplementary Dataset. S4A, filter column BE “Transport”). In seeds, VDAC isoforms and several protein translocators were significantly elevated, whereas a range of substrate carriers predominated in leaves.

To pin down the major differences in mitochondrial energy metabolism between the two tissues we focused on the quantitative distribution of dehydrogenases since these enzymes directly provide the electrons to fuel oxidative phosphorylation (Fig. 5, Supplementary Dataset S4A, filter column BI, Supplementary Dataset S4D). Formate dehydrogenase was dominant in seed mitochondria contributing ∼3% of total mitochondrial protein mass compared to only ∼0.3% in leaves. The pyruvate dehydrogenase complex and malate dehydrogenase and the glycine cleavage system were also high abundant but not as prominent in seeds as in leaves. The oxoglutarate dehydrogenase complex, several dehydrogenases involved in amino acid catabolism as well as an uncharacterized alcohol dehydrogenase were increased in seed mitochondria whereas isocitrate dehydrogenase was more abundant in the leaves.

**Fig. 5:**
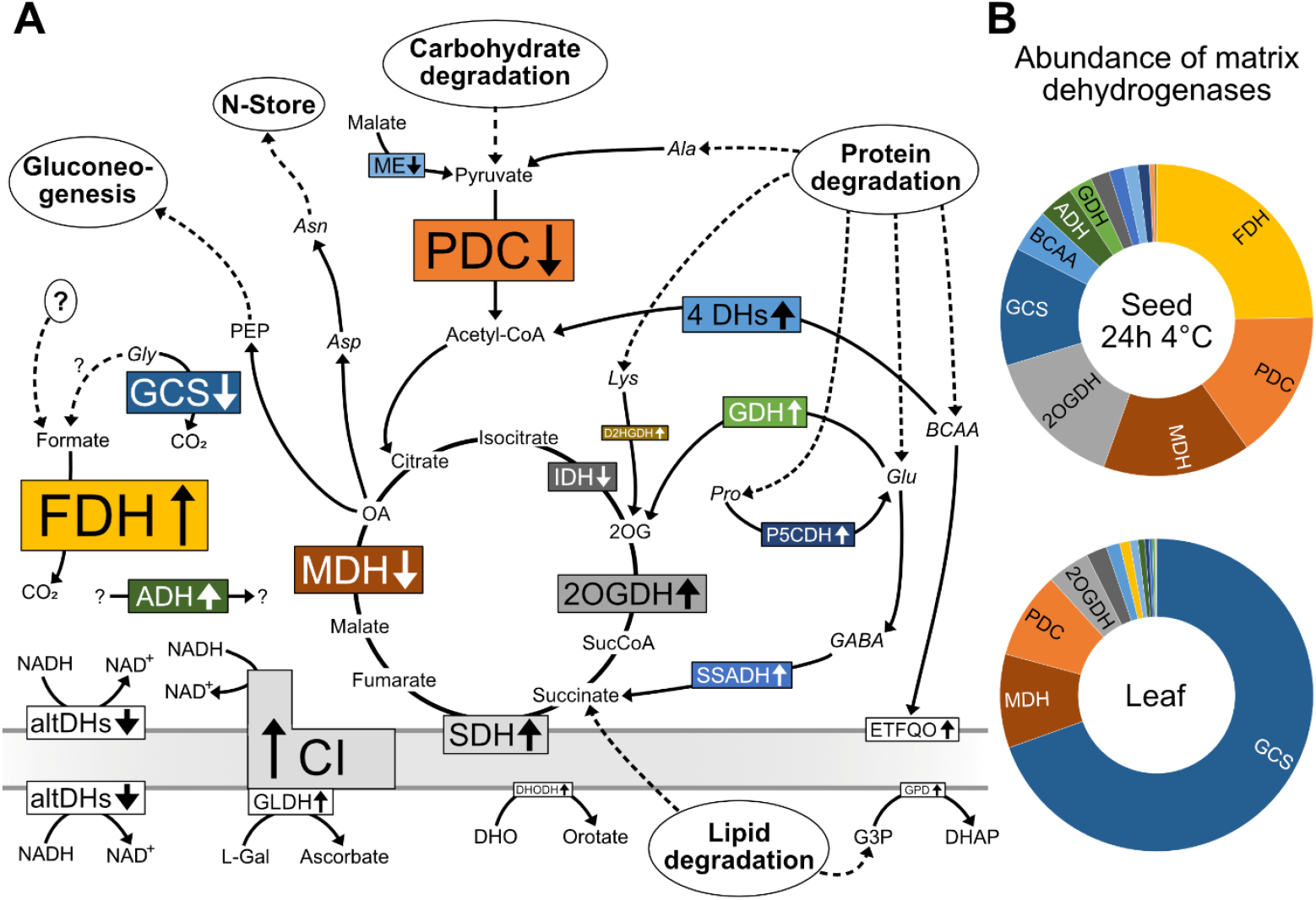
Dehydrogenases in seed mitochondrial energy metabolism. Schematic representation of mitochondrial metabolic pathways, highlighting the roles of dehydrogenases and their associated pathways in seeds. (**A**) The text size corresponds to the total abundance of the individual dehydrogenases or dehydrogenase complexes in quiescent seed mitochondria (iBAQ%), which is also illustrated in the respective diagram on the right. Arrows indicate relative protein abundance in seed vs. leaf mitochondria. Remobilization of seed storage compounds provides substrates for mitochondrial energy metabolism and gluconeogenesis (dashed lines). Dehydrogenases: ADH: alcohol dehydrogenase; altDH: alternative NADH dehydrogenases; BCAA/4 DHs include the four dehydrogenases involved in branched-chain amino acid catabolism (branched-chain alpha-keto acid dehydrogenase complex, isovaleryl-CoA-dehydrogenase, 3-hydroxyisobutyrate dehydrogenase, methylmalonate-semialdehyde dehydrogenase); D2HGDH: D-2-hydroxyglutarate dehydrogenase; DHODH: dihydroorotate dehydrogenase; ETFQO: electron-transfer flavoprotein:ubiquinone oxidoreductase; FDH: formate dehydrogenase; GCS: glycine cleavage system; GDH: glutamate dehydrogenase; GLDH: L-galactono-1,4-lactone dehydrogenase; GPD: glycerol-3-phosphate dehydrogenase; IDH: isocitrate dehydrogenase; MDH: malate dehydrogenase; ME: malic enzyme; P5CDH: delta-1-pyrroline-5-carboxylate dehydrogenase; PDC: pyruvate deyhydrogenase complex; SDH: respiratory chain complex II (succinate dehydrogenase); SSADH: succinic semialdehyde dehydrogenase; 2OGDH: 2-oxoglutarate dehydrogenase complex. Proline dehydrogenase and D-Lactate dehydrogenase were not detected in our samples. DHAP: dihydroxyacetone phosphate; DHO: dihydroorotate; G3P: glycerol-3-phosphate; L-Gal: L-galactono-1,4-lactone; OA: Oxaloacetate; PEP: Phosphoenolpyruvate; SucCoA: Succinyl-CoA; 2OG: 2-Oxoglutarate. Abundance of matrix dehydrogenases in mitochondria of seeds imbibed for 24h at 4°C and leaf based on their mass fraction (%iBAQ). The complete dataset is provided as Supplementary Dataset S4.

Seed mitochondria were further distinguished by a larger fraction of uncharacterized proteins, accounting for 12% of total mitochondrial protein mass compared with 5% in leaves (Fig. 4A). Approximately one quarter of the proteins with significantly higher abundance in seeds remain functionally uncharacterized (Fig. 4B, blue dots, Supplementary Dataset S4A, filter column BF). Among them, 32 belonged to the 20% most abundant proteins in seed mitochondria (Fig. 4B, dark blue dots) and 64 proteins showed only week homology to *A. thaliana* and might thus have lupin- or legume-specific functions. Coexpression analysis suggests some of these uncharacterized proteins might be involved in RNA processing, OXPHOS, or transport processes (Supplementary Dataset S5). Moreover, 43 of the newly identified mitochondrial protein candidates were significantly more abundant in seeds than leaves and thus might have seed-specific functions.

## Discussion

Successful germination critically depends on the rapid re-establishment of energy metabolism in seeds, with mitochondria playing a central role as the primary source of ATP supply. In this study, we provide the first comprehensive proteomic characterization of mitochondria isolated from *Lupinus albus* seeds, thereby expanding the understanding of mitochondrial function during the onset of germination. Our findings reveal both conserved features and novel aspects of mitochondrial metabolism in a legume with protein-rich reserves.

### Seed mitochondria are preconfigured for metabolic activation

Our shotgun proteomic and BN/SDS-PAGE analyses demonstrate that seed mitochondria contain fully assembled OXPHOS complexes and supercomplexes even in the quiescent state. This is consistent with recent findings in *Arabidopsis thaliana*, where cristae structures and intact electron transport chain complexes were observed in dry seeds, indicating that mitochondria are primed to support energy production immediately upon imbibition (Ditz et al., 2025). The ability to rapidly resume respiration ensures efficient ATP production during the early germination process, which is tightly linked to seed vigor (Ferguson et al., 1990; Kühn et al., 2015; Racca et al., 2022). Our quantitative proteome confirms this preconfiguration, with ∼40% of protein mass dedicated to OXPHOS, highlighting the centrality of respiration for energy supply in seeds during early germination.

### Diversification of respiratory substrates in seed mitochondria

Comparison of the lupin seed mitochondrial proteome with that of leaf mitochondria reveals striking differences that reflect the contrasting metabolic demands of these tissues. Whereas leaf mitochondria integrate into photorespiration and cooperate with chloroplast metabolism, seed mitochondria operate in a photosynthetically inactive environment, where their primary function is to support the mobilization of storage reserves and to generate ATP for biosynthetic processes. This is consistent with the observation that respiratory chain (MRC) components are highly abundant and fully assembled in dry seeds, whereas TCA cycle proteins are less prominent. Such a profile indicates reliance on alternative sources of reducing equivalents rather than a canonical TCA cycle.

The quantitative profile of dehydrogenases provides insights into how seed mitochondria sustain respiration independently of a full TCA cycle. A remarkable feature of lupin seed mitochondria is the exceptionally high abundance of NAD-dependent formate dehydrogenase (FDH), which ranks third among all identified mitochondrial proteins. FDH catalyzes the oxidation of formate to CO_2_ with concomitant production of NADH, thereby directly linking detoxification with energy supply. Previous studies demonstrated that FDH is highly abundant in mitochondria of non-photosynthetic tissues such as potato tubers, dark-grown shoots, storage roots, and cell suspensions, but much less present in leaf mitochondria (Des Francs-Small et al., 1992). Its abundance can reach up to 9% of mitochondrial protein in tubers, with activities 5–8 times higher than in leaves, suggesting a key metabolic role in non-green tissues (Des Francs-Small et al., 1993). High activities have also been observed in mature seeds (Davison, 1949). Our findings confirm that FDH is a key metabolic enzyme in seed mitochondria and suggest that formate detoxification may be particularly important during early germination. The metabolic sources of formate in seeds remain incompletely understood. Potential pathways include pectin demethylation, glyoxylate metabolism, and amino acid degradation, all of which are active during reserve mobilization in seeds (Igamberdiev et al.,1999). In this context, the high activity of FDH may serve as an adaptive mechanism to prevent formate accumulation while simultaneously fueling respiration during the metabolic transition.

The observed differences in quantitative dehydrogenase composition between mitochondria from lupin seeds and leaves as well as proteome dynamics during seed transition from a dormant to a metabolically active state reflect adaptations to remobilization of the seed storage compounds and are well in line with a combined metabolism of fatty acids and amino acids postulated based on previous findings (Angermann et al., 2024). The dehydrogenases involved in mitochondrial amino acid catabolic pathways are of higher abundance in seed compared to leaf mitochondria and thus potentially contribute more to ATP production. In addition, amino acid degradation pathways are induced during early germination. Ample availability of glycine would promote transamination to glyoxylate and support a linear flux mode through parts of the glyoxylate cycle being combined with acetyl-CoA from fatty acid β-oxidation at the level of malate synthase to produce oxaloacetate for gluconeogenesis (Angermann et al., 2024). Malate dehydrogenase, catalyzing the oxidation step of malate to oxaloacetate in this pathway, is much more abundant in lupin compared to Arabidopsis seed mitochondria (1.7% vs. 0.15% of the mitoproteome, *A. thaliana* data from Ditz et al. (2025)) and it increases during germination.

### Coenzyme metabolism and mitochondrial activation during early germination

Our quantitative proteome data also reveal that coenzyme metabolism is upregulated in *Lupinus albus* seed mitochondria during the early stages of germination, with elevated levels of enzymes involved in lipoic acid and thiamine biosynthesis as well as Fe–S cluster assembly. These cofactors are essential for the activity of mitochondrial dehydrogenase complexes, including pyruvate dehydrogenase, 2-oxoglutarate dehydrogenase, and glycine decarboxylase. Disruption of lipoic acid biosynthesis causes embryo lethality in Arabidopsis (Ewald et al., 2014), while thiamine deficiency severely impairs seedling establishment (Raschke et al., 2007). Fe–S clusters are indispensable for respiratory chain function and mitochondrial metabolism more broadly (Lill and Freibert, 2020), and defects in their assembly compromise embryo development as well as seedling establishment and root growth (Bernard et al., 2009; Moseler et al., 2015). The simultaneous induction of these pathways suggests that the activation of mitochondrial metabolism during germination requires coordinated cofactor provisioning to ensure that enzyme complexes are fully functional.

### Transporter complement reflects energetic and metabolic priorities of seed mitochondria

In addition to metabolic enzymes, the distinct profile of mitochondrial transporters highlights functional specialization in seeds. The high abundance of the ADP/ATP carrier (ANT) underscores the central role of mitochondria in ATP provision to the cytosol during germination, in contrast to the more balanced role of ANT in leaf mitochondria, where photorespiratory metabolism predominates. Similarly, the very high levels of voltage-dependent anion channel (VDAC) in seed mitochondria suggest a strong demand for metabolite exchange across the outer membrane during the initiation of metabolism. Such elevated transport capacity is consistent with the requirement for efficient coordination between storage compound degradation, respiratory activity, and biosynthesis in germinating seeds. VDAC has also been identified as the protein with the highest copy number in mitochondria isolated from a heterotrophic *Arabidopsis thaliana* cell suspension culture (Fuchs et al. 2020). A large fraction of the high abundant seed mitochondrial metabolite transporters has not been characterized yet and might include functional adaptations to storage compound metabolism.

### Lupin seed mitochondria reveal potential legume-specific functions

Beyond the conserved features shared with Arabidopsis, the lupin seed mitochondrial proteome exhibits specific signatures that may reflect legume-specific adaptations. Two regulatory proteins, the cystationine beta-synthase family protein CBSX3 (A0A6A5NX76/AT5G10860) and EARLY NODULIN 93 (ENOD93, A0A6A5MA33/AT5G25940), are among the most abundant proteins in lupin seed mitochondria but are much less prominent in Arabidopsis. CBSX3, a regulator of thioredoxin-o, is implicated in redox homeostasis and may be critical in balancing ROS production during the early burst of respiratory activity (Yoo et al., 2011; Shin et al., 2020; Ok et al., 2012). ENOD93, originally described in the context of legume nodulation, has recently been identified as a regulator of cytochrome c oxidase activity (Lee et al., 2024). By modulating respiratory ATP production, ENOD93 maintains mitochondrial energy balance, and its loss leads to reduced ADP-dependent respiration and altered membrane potential. Its high abundance in lupin seed compared to leaf mitochondria indicates that additional layers of respiratory regulation fine-tune energy balance during germination.

Our study also revealed a high proportion of uncharacterized proteins in lupin seed mitochondria, accounting for ∼12% of total protein mass. Many of these proteins lack close homologs in Arabidopsis, raising the possibility of legume-specific mitochondrial adaptations. Some uncharacterized proteins were among the most abundant components of the seed mitochondrial proteome and showed dynamic changes during germination, suggesting important but as yet undefined roles. Coexpression analyses linked subsets of these proteins to mitochondrial translation, ribosome biogenesis, and cofactor biosynthesis, highlighting them as promising candidates for functional studies.

### Lupin as a model system for seed mitochondrial biology

Our study demonstrates the high potential of lupin seeds as a model to study the role of mitochondria during germination. Their large size allows for efficient organelle purification and high protein yields without the need for elaborate tag-based enrichment procedures. Our curated annotation database provides the opportunity to use the established tools and extensive resources available for *Arabidopsis thaliana*. In total, we identified 1,162 seed mitochondrial proteins in *L. albus*, closely comparable to a recent Arabidopsis seed mitochondrial proteome dataset comprising 1,148 distinct proteins derived from an in-depth analysis of the total seed proteome (Ditz et al., 2025). The overlap of 749 distinct proteins between the two datasets underscores a strong conservation of mitochondrial function across species, while the observed differences open perspectives to investigate species-specific adaptations.

### Conclusion

Our study provides a comprehensive analysis of the lupin seed mitochondrial proteome, revealing both conserved features shared with Arabidopsis and unique aspects likely reflecting legume-specific adaptations. Seed mitochondria are primed with fully assembled OXPHOS complexes, enriched in specialized dehydrogenases and transporters, and undergo targeted remodeling of coenzyme metabolism and substrate utilization pathways during germination. The large fraction of abundant, uncharacterized proteins highlights the potential for discovering new mitochondrial functions specific to seed biology and legumes. Together, these findings establish white lupin as a valuable model for studying the mitochondrial basis of seed vigor and germination.

## Materials and Methods

### Plant material

White lupin (*Lupinus albus* cv. “Nelly”) seeds were obtained from Revierberatung Wolmersdorf GmbH & Co. KG, Wolmersdorf, Germany (Order number: 34400) and stored in the dark at 4°C until use. Seed samples were cultivated as follow: 0h seeds were used for protein extraction or mitochondrial isolation directly without any treatment. For imbibition seeds were soaked for 24h in demineralized water at 4°C or 22°C (RT). Fresh plant material was used to isolate mitochondria (five pools of 100 g (fresh weight)) or measure respiration rates (single seeds).

For total protein extraction five pools of 10 seeds were used. The seed coats were removed and the remaining plant organs were deep-frozen in liquid nitrogen. The plant material was lyophilized in an Alpha 1-2 LD+ freeze dryer (Christ, Osterode, Germany). The dried material was ground into powder.

Leaf mitochondria were isolated from 28-day old plants. For plant cultivation, seeds were soaked in demineralized water at 22°C for 16h and then transferred to water-saturated expanded clay substrate (LamstedtDan, 4 – 8 mm, Fibo ExClay Deutschland GmbH, Lamstedt, Germany). Plants were cultivated in a phyto chamber (22 – 24°C, 16h light, 8h dark, 110 μmol · m^−2^ · s^−1^ light). At day 28 leaves of several plants were collected and pooled to 5 batches of 100 g (fresh weight) and used for isolation of mitochondria.

The germination capacity of seeds that had been soaked for 24 hours at either 4°C or RT was tested. This was achieved by placing 100 seeds from each group on a wet paper towel in the dark at room temperature and allowing them to germinate. 99 of the RT-soaked seeds germinated within 6 hours. Germination of the seeds that had been imbibed at 4°C for 24h was delayed. Three days after placing them on the paper towel, a total of 98 seeds had germinated.

### Mitochondrial preparation

Mitochondria were isolated from *Lupinus albus* seeds following the protocol described in Werhahn et al. (2001). All steps are prepared in a cold room or on ice. In short, 100 g (fresh weight) of each sample were homogenized in pre-cooled extraction buffer (450 mM sucrose, 15 mM 3-(N-morpholino)-propanesulfonic acid (MOPS), 1.5 mM EGTA, 0.6% (w/v) polyvinylpyrrolidone 40, 0.2% (w/v) bovine serum albumin, 10 mM sodium ascorbate, 10 mM cysteine, 0.2 mM phenylmethylsulfonyl fluoride (PMSF), pH 7.4) by a blender and filtered through two layers of muslin. Filtrate was centrifuged for 5 minutes at 4°C and 2700 × g. The supernatant was centrifuged for 5 minutes at 4°C and 8300 × g. Mitochondria were pelletized by centrifugation (10 minutes at 4°C and 17 000 × g). The mitochondrial pellet was resuspended in wash buffer (300 mM sucrose, 10 mM MOPS, 1 mM EGTA, 0.2 mM PMSF, pH 7.2) and layered on a three-step Percoll gradient (18% [v/v], 23% [v/v], and 40% [v/v] Percoll in 300 mM sucrose and 10 mM MOPS, pH 7.2). After centrifugation for 90 minutes at 4°C at 70 000 × g the mitochondria can be isolated from the 23% / 40% interphase. The purified mitochondria were washed twice in resuspension buffer (400 mM mannitol, 1 mM EGTA, 0.2 mM PMSF, 10 mM Tricine, pH 7.2) and centrifuged for 10 minutes at 4°C at 14 500 × g. Isolated mitochondria were snap-frozen in liquid nitrogen and stored at −80°C.

Mitochondria from 28-day old leaves were purified by differential centrifugation and Percoll density gradient centrifugation following the protocol by Keech et al. (2005). All steps were performed in a cold room or on ice. In short, Percoll gradients were prepared as follows: 50% (v/v) Percoll and 50% (v/v) buffer (600 mM sucrose, 20 mM TES, 2 mM EDTA, 20 mM KH2PO4, 2 mM glycine, pH 7.5), centrifuged for 40 minutes at 4°C at 69 400 × g and stored at 4°C until use. 100 g (fresh weight) of each sample were homogenized in pre-cooled extraction buffer (300 mM sucrose, 60 mM N-tris [hydroxymethyl]-methyl-2-aminoethanesulphonic acid (TES), 25 mM sodium pyrophosphate decahydrate, 10 mM KH2PO4, 2 mM EDTA, 1 mM glycine, 1% polyvinylpyrrolidone 40, 1% (w/v) bovine serum albumin, 50 mM sodium ascorbate, 20 mM cysteine, pH 8) in a blender and filtered through two layers of muslin. The filtrate was centrifuged for 5 minutes at 4°C and 2 500 × g. The supernatant was centrifuged for 15 minutes at 4°C and 15 100 × g. The mitochondrial pellet was resuspended in wash buffer (300 mM sucrose, 10 mM TES, 10 mM KH2PO4, pH 7.5) and layered on a prepared Percoll gradient. After centrifugation for 20 minutes at 4°C at 17 400 × g the mitochondria formed a whitish band close to the bottom of the tube. This band was aspirated, washed twice in wash buffer and centrifuged for 20 minutes at 4°C at 17 200 × g. Isolated mitochondria were snap-frozen in liquid nitrogen and stored at −80°C.

### Respiration measurements

Respiration rates were measured using two seeds per replicate in 6 ml demineralized water under ambient conditions (22°C) or at 4°C (in a fridge) using an Oxygraph+ with DW3 electrode chamber (Hansatech Instruments Ltd, Norfolk, UK).

### 2D Blue native (BN)/SDS polyacrylamide gel electrophoresis (PAGE)

The protein content of the isolated mitochondrial samples was quantified according to Angermann et al. (2024). In short, ice-cold methanol (100%) was added to the isolated mitochondria and incubated for 20 minutes at −20°C. After centrifugation (5 minutes, 4°C, 14 200 × g) the pellet was dissolved in 200 ml of 10 mM NaOH containing 0.2% SDS (v/w) and incubated for 5 minutes at 60°C, shaking. After centrifugation (10 minutes, RT, 7 000 × g), the protein content was quantified by the Pierce BCA Protein Assay Kit (Thermo Fisher Scientific, Rockford, Illinois, USA). Globulin was used as a standard.

2D BN/SDS PAGE was performed according to Wittig et al. (2006). In summary, 1D BN PAGE was used to analyse mitochondrial protein complexes. A gradient gel (16 × 16 × 0.15 cm) was created using a MX40 gradient mixer, with two chambers filled with the following solutions: Front chamber: Gel buffer BN (250 mM ACA, 25 mM Bis-Tris, pH 7.0) containing 4.5% (v/v) acrylamide. Back chamber: Gel buffer BN containing 16% (v/v) acrylamide and 19% (v/v) glycerol. Once the gradient gel had polymerised, a sample gel (gel buffer BN, 4% acrylamide) was added on top. The wash buffer was removed from the mitochondrial samples by centrifugation and the proteins were extracted from the organelles using a solubilisation buffer (30 mM HEPES, 150 mM potassium acetate, 10% glycerol and 5% digitonin, pH 7.4) at a ratio of 100 μl per 10 μg of mitochondrial protein. Following a 10-minute incubation on ice, the membrane aggregates and other insoluble materials were removed by centrifugation at 18 300 × g for 10 minutes. Subsequently, Coomassie Blue was added to the sample to achieve a final concentration of 1% (w/v). Blue native PAGE was conducted using the following buffers: Anode buffer: 50 mM Bis-Tris, pH 7.0. Cathode buffer: 50 mM tricine, 15 mM Bis-Tris and 0.02% Coomassie, pH 7.0. Two steps were involved in performing the electrophoresis: i) max. 100 V (current set to 15 mA) for 45 minutes; and ii) max. 15 mA with a voltage of 500 V for 11 hours. The gel was then fixed and subjected to Coomassie colloidal staining (Neuhoff et al., 1988) (fixation: 15% ethanol and 10% acetic acid with a 2-hour incubation, followed by staining with 5% (w/v) Coomassie in 2% (w/v) ortho phosphoric acid and 10% (w/v) ammonia sulfate with an overnight incubation) or utilised for a second dimension SDS-PAGE.

SDS-PAGE (second gel dimension after BN PAGE): A lane was removed from the blue native gel and treated with a solution of 1% SDS and 1% beta-mercaptoethanol for half an hour. Then, the lane was submerged in water for 30 seconds. The gel lane was then placed between two glass plates. A separation gel containing 16% (v/v) acrylamide and 12% (v/v) glycerol in SDS gel buffer (1 M Tris, 0.1% SDS, pH 8.5) and a spacer gel containing 10% (v/v) acrylamide in SDS gel buffer were poured directly on top of each other between the plates. After polymerisation, a sample gel containing 10% (v/v) acrylamide, 10% (v/v) glycerol and 0.1% (w/v) SDS in BN gel buffer was poured between the SDS gel and the blue native gel strip. Finally, the gel (19 × 16 × 0.1 cm) was placed in the chamber and anode buffer (0.2 M Tris, pH 8.9) and cathode buffer (0.1 M Tris, 0.1 M Tricine and 0.1% SDS, pH 8.5) were added. Gel electrophoresis was then performed for approximately eighteen hours at 30 mA (maximum voltage: 500 V) and stopped when the blue running front entered the anode buffer. The gel was then fixed and stained using the Coomassie colloidal method, as previously described.

### Protein extraction, digestion and sample preparation for shotgun proteome analysis via mass spectrometry

The solvent precipitation, single-pot, solid-phase-enhanced sample preparation (SP4) protocol was used for mass spectrometry sample preparation as described in Heinemann et al. (2025). In brief, the proteins were solubilised and denatured by incubation at 60°C for 30 minutes in SDT buffer (4% SDS, 0.1 M DTT, 0.1 M Tris, pH 7.6) (Mikulášek et al., 2021). The samples were then sonicated and centrifuged at 20 000 × g for 10 minutes. Then, 30 μl of the extract was transferred to a new tube and mixed with 7.5 μl of iodoacetamide (IAM, 0.1 M). The mixture was incubated for 30 minutes in the dark to alkylate the reduced disulfide bridges. Then, 2 μl of DTT (0.1 M) was added to neutralise any excess IAM.

The preparation of the glass beads/ACN suspension, the protein precipitation and the washing steps were performed as described by Johnston et al. (2022). Approximately 400 μg of glass beads were used per sample (approximately 30 μg of protein). The purified proteins were digested with 0.5 μg of mass spectrometry-grade trypsin (Promega) for 16 hours at 37°C with shaking. The peptide-containing supernatant was collected in a low-binding tube and acidified with 1 μl of formic acid. The peptides were desalted using 50 mg Sep-Pak C18 columns (WAT054960, Waters, Milford, MA, USA) and quantified with the Pierce Quantitative Colourimetric Peptide Assay Kit (Thermo Fisher Scientific, Rockford, IL, USA). The samples were finally diluted to a concentration of 400 ng μl^-1^ in 0.1% FA.

### Quantitative shotgun proteomics by ion Mobility Mass Spectrometry (IMS-MS/MS)

Four hundred nanograms of peptides were injected via a nanoElute2 UHPLC (Bruker Daltonics GmbH, Bremen, Germany) and separated on an analytical reversed-phase C18 column (Aurora Ultimate 25 cm × 75 µm, 1.6 µm, 120 Å; IonOpticks). Using MS grade water and a multi-staged acetonitrile gradient containing 0.1% formic acid (0 min, 2%; 54 min, 25%; 60 min, 37%; 62 min, 95%; 70 min, 95%), peptides were eluted and ionized by electrospray ionization with a CaptiveSpray2 ion source and analyzed on a timsTOF-HT mass spectrometer (Bruker Daltonics GmbH, Bremen, Germany), with the following DDA-PASEF settings: Ion mobility window of 0.7 – 1.5 cm^-2^ V^-1^ s^-1^, 4 PASEF ramps, target intensity 14 500 (threshold 1 200), and a cycle time of ∼0.53 s.

### Data processing and functional annotation

The ion mobility spectrometry (IMS)–MS/MS spectra from all experiments were analyzed with the MaxQuant software (Cox and Mann, 2008) using default search parameters and the proteome database of *Lupinus albus* published by Xu et al. (2020) on UniProt.org (UP000464885) for protein identification. The calculation of label-free quantification (LFQ) values and intensity-based absolute quantification (iBAQ) values were both enabled. Data evaluation was performed using Perseus (Tyanova et al., 2016). Proteins were excluded from further analysis if they were not detected in at least two out of three or four out of five biological replicates in at least one of the sample groups. Subsequently, missing values were replaced with randomly chosen low values from a normal distribution. Significant changes were calculated using Student’s t-tests (p = 0.05). For comparisons between mitochondrial proteomes datasets were filtered for mitochondrial localization to remove contaminations from other compartments and iBAQ values were normalized prior to statistical evaluation. A recently published annotation database of *Lupinus albus* was used to obtain further information such as subcellular localization and metabolic pathway involvement (Angermann et al., 2024). Fisher’s exact tests were performed in Perseus to identify significantly enriched or depleted metabolic pathways within the mitochondrial-localised proteins.

## Author contributions

CA and TMH designed the research; CA performed and evaluated all experiments; CA, HPB, and TMH analyzed the data; CA and TMH wrote the manuscript with support from HPB; TMH agrees to serve as the author responsible for contact and ensures communication.

## Data availability

The mass spectrometry proteomics data have been deposited to the ProteomeXchange Consortium (http://proteomecentral.proteomexchange.org) via the PRIDE partner repository (Perez-Riverol et al. 2022) with the Dataset identifier PXD068370.

## Supplementary Data

Supplementary Dataset S1: Complete proteomics datasets of total seed and mitochondrial preparations

Supplementary Dataset S2: Identification of new mitochondrial proteins

Supplementary Dataset S3: Coexpression analysis of new mitochondrial proteins

Supplementary Dataset S4: *Lupinus albus* seed and leaf mitochondrial proteomes

Supplementary Dataset S5: Coexpression analysis of uncharacterized seed mitochondrial proteins

Supplementary Figure S1: Specification of thresholds for identifying new mitochondrial proteins

## Acknowledgments

We would like to thank Björn Heinemann for his invaluable help with organizing, conducting and evaluating the experiments. We thank Dagmar Lewejohann for cultivation of leaf material, isolation of leaf mitochondrial fractions and skillful preparation of the gels. We thank Christina Mack for skillful technical assistance.

## Supplementary Figures

**Supplementary Figure S1:**
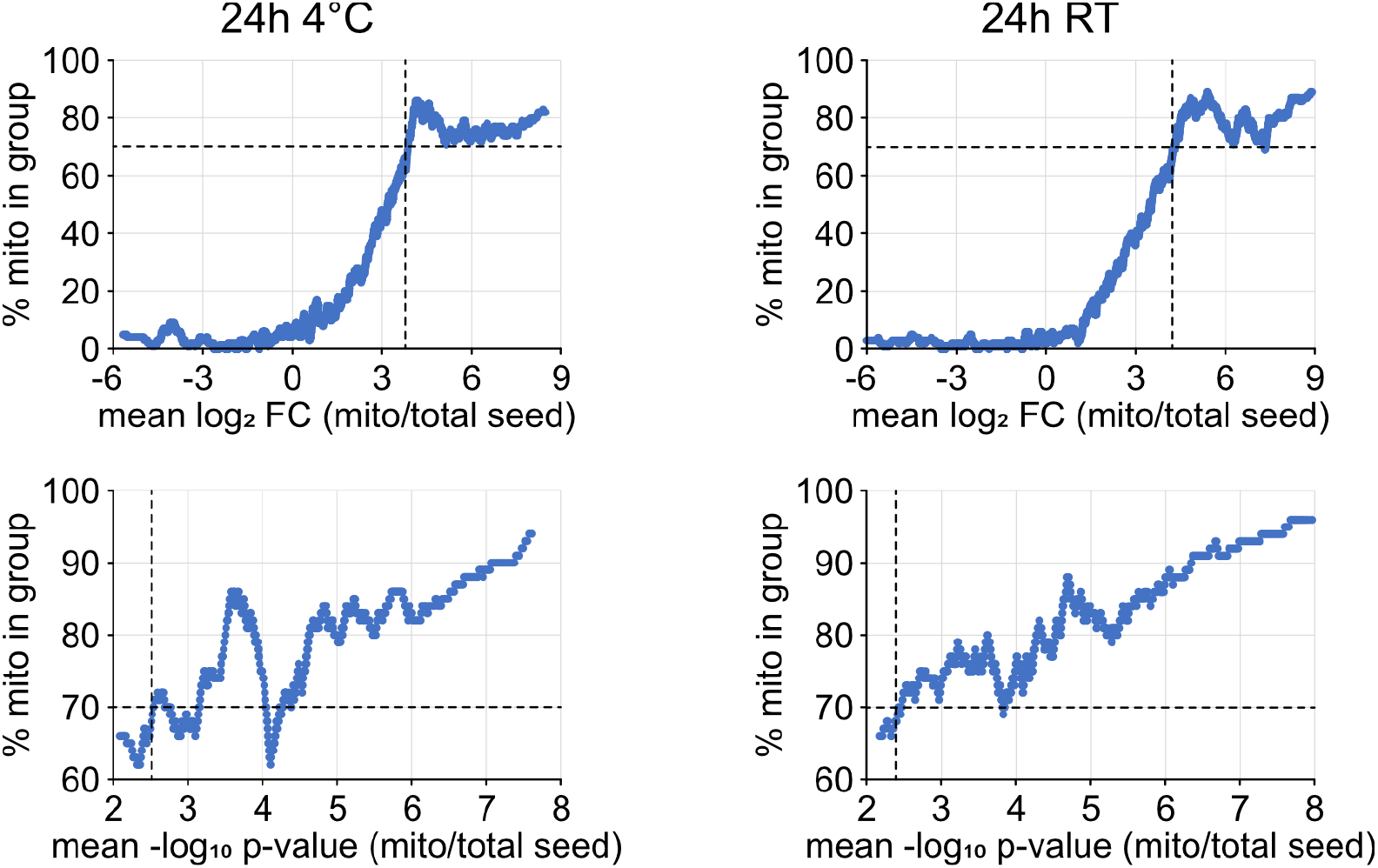
Specification of thresholds for identifying new mitochondrial proteins. The moving average of the enrichment factor (mean log_2_-ratio of protein abundances in mitochondrial fractions and whole seed proteomes) and the respective −log_10_ p-values was calculated for a window size of 100 proteins and plotted against the percentage of proteins in the same group with a mitochondrial annotation in SUBA5. The threshold for identifying new mitochondrial protein candidates was set at 70% mitochondrial annotations (dotted lines).

